# Discovery and dynamic pharmacology of µ-opioid receptor positive allosteric modulators

**DOI:** 10.64898/2026.02.20.707058

**Authors:** Evan S. O’Brien, Junzheng Wang, Parthasaradhireddy Tanguturi, Mengchu Li, Elizabeth White, Yuki Shiimura, Barnali Paul, Kevin Appourchaux, Kaavya Krishna Kumar, Weijiao Huang, Susruta Majumdar, John R. Traynor, John M. Streicher, Chunlai Chen, Brian K. Kobilka

## Abstract

Opioid agonists such as morphine and fentanyl exert analgesic effects by binding and activating the µ-opioid receptor (µOR), yet agonism of the µOR causes a slate of serious side effects. µOR-mediated addiction and respiratory depression are the major causes of the current opioid overdose crisis, largely driven by the explosion in illicit use of fentanyl, a potent opioid receptor full agonist. Given these serious side effects (and high resulting societal cost), molecules that act as analgesics with distinct mechanisms of action are of great interest. Positive allosteric modulators (PAMs) of the µOR have the potential to avoid many off-target side effects of conventional opioid orthosteric agonists by enhancing the signaling properties of natural opioid peptide systems. We used a DNA-encoded chemical library screening approach to selectively discover active-state-specific µOR PAMs. Two out of 3 selected prospective PAMs displayed the anticipated enhancement in agonist activity. The most effective of these compounds enhanced the activity of all orthosteric opioid agonists tested, including the native opioid peptide met-enkephalin. Little is known about the underlying dynamic basis of allosteric modulation of Family A GPCRs like the µOR. To that end, we used single-molecule fluorescence resonance energy transfer experiments to detail the impact that our novel µOR PAM has on the dynamic activation behavior of a key region on the intracellular face of the receptor. Our results here provide both a new chemical scaffold that acts as a µOR PAM and detailed pharmacological and dynamic insights into its mechanism of action.

## Introduction

Opioids derived from the poppy plant *Papaver somniferum* have been used extensively, both recreationally and as powerful pain relievers (e.g. heroin). Their semi-synthetic relatives codeine and oxycodone are also analgesics that are widely used clinically (along with the fully synthetic opioid fentanyl) despite their known potential for addiction, abuse, and respiratory depression. Opioid drugs generally are considered front-line therapeutics for managing severe acute pain but have historically been prescribed for longer-term chronic pain as well; this longer-term use of particularly oxycodone has been a main driver of the current opioid overdose crisis. Concurrent with this rise in over-prescribed long-term pain management opioids, fully synthetic opioids like fentanyl have exploded as cheap recreational alternatives to opioids like oxycodone or even heroin.

All these opioids work principally by binding to the same site (**Fig. S1a**) as endogenous opioid peptides such as the enkephalins and endorphins on the µ-opioid receptor (µOR), though all have different affinities for the receptor and various extents of signaling (efficacies). Given the extensive drawbacks of current opioids in long-term settings, there is a pressing need for new drugs that act as analgesics with distinct mechanisms of action, including activation of other GPCRs that are involved in pain regulation^1–3^. It may also be possible to take advantage of the extensive analgesia mediated by the µOR with molecules that bind at sites distinct from the conventional orthosteric site and activate the receptor. Such positive allosteric modulators (PAMs) may be more selective for the µOR over other opioid receptor family members κ-opioid receptor (κOR) and δ-opioid receptor (δOR). Past receptor selectivity, PAMs have the potential to provide more selective therapeutic effects by working synergistically with the body’s endogenous opioid systems to enhance natural analgesic effects in a spatially and temporally regulated manner, without activating unwanted reward and respiratory depression circuits^4^. The so-called “ceiling effects” of allosteric modulators, dependent on the cooperativity between the allosteric site and orthosteric site, can also result in a diminished propensity for µOR-induced overdose^5^. In certain situations, allosteric modulators display “probe dependence”, where they are able to selectively enhance the activity of particular orthosteric molecules. For instance, PAMs of the µOR that selectively enhance endogenous peptide agonism rather than that from morphine or fentanyl would be highly desirable, as would negative allosteric modulators (NAMs) that are less effective against endogenous agonists and could be used to help manage opioid use disorder.

Several µOR PAM chemical series have been discovered and developed, including the BMS^6,7^ and MS^8^ compounds and their derivatives^9^, some of which show promising therapeutic activities as adjuncts to opioid activities *in vivo*^9^. However, the potencies of these compounds remain low, requiring high doses, which are also complicated by their relatively poor solubilities. New chemical series of PAMs, as well as clearly defined mechanisms of action, are therefore of great potential utility.

The discovery of such allosteric modulators can be streamlined through a directed screen of large DNA-encoded chemical libraries (DELs) (**Fig. 1a**). DELs can be composed of billions or even trillions of molecules, all of which are covalently attached to a readily sequenced DNA barcode^10–12^. We have previously demonstrated the utility of this approach in the discovery and characterization of a specific µOR NAM that works *in vivo* to cooperatively “boost” the activity of the key opioid overdose reversal compound naloxone^13^. Previous work using DELs has successfully identified β_2_ adrenergic receptor PAMs and NAMs^14–17^. We used a screening strategy that can be used more broadly to identify PAM and NAM compounds for any receptor of interest by “steering” the hits towards sites distinct from the receptor orthosteric site and ensuring that any selected compounds are likely to bind to and stabilize the desired active or inactive state ensembles (**Fig. 1**). This approach resulted in the efficient discovery of new µOR PAM chemical scaffolds, the best of which broadly enhanced agonist-induced activity at the receptor while acting as nearly a full-agonist in the absence of orthosteric compound (i.e. ago-PAM activity). Using single-molecule fluorescence resonance energy transfer (smFRET) experiments, we then show that the PAM enhances the population of an agonist-associated intermediate state of the receptor, and that this increased activation directly leads to increased G-protein association. Such PAM molecules will hopefully serve as new chemical material for next-generation analgesic compounds capable of “boosting” our body’s natural pain response systems without being addictive or causing overdoses.

**Figure 1.**
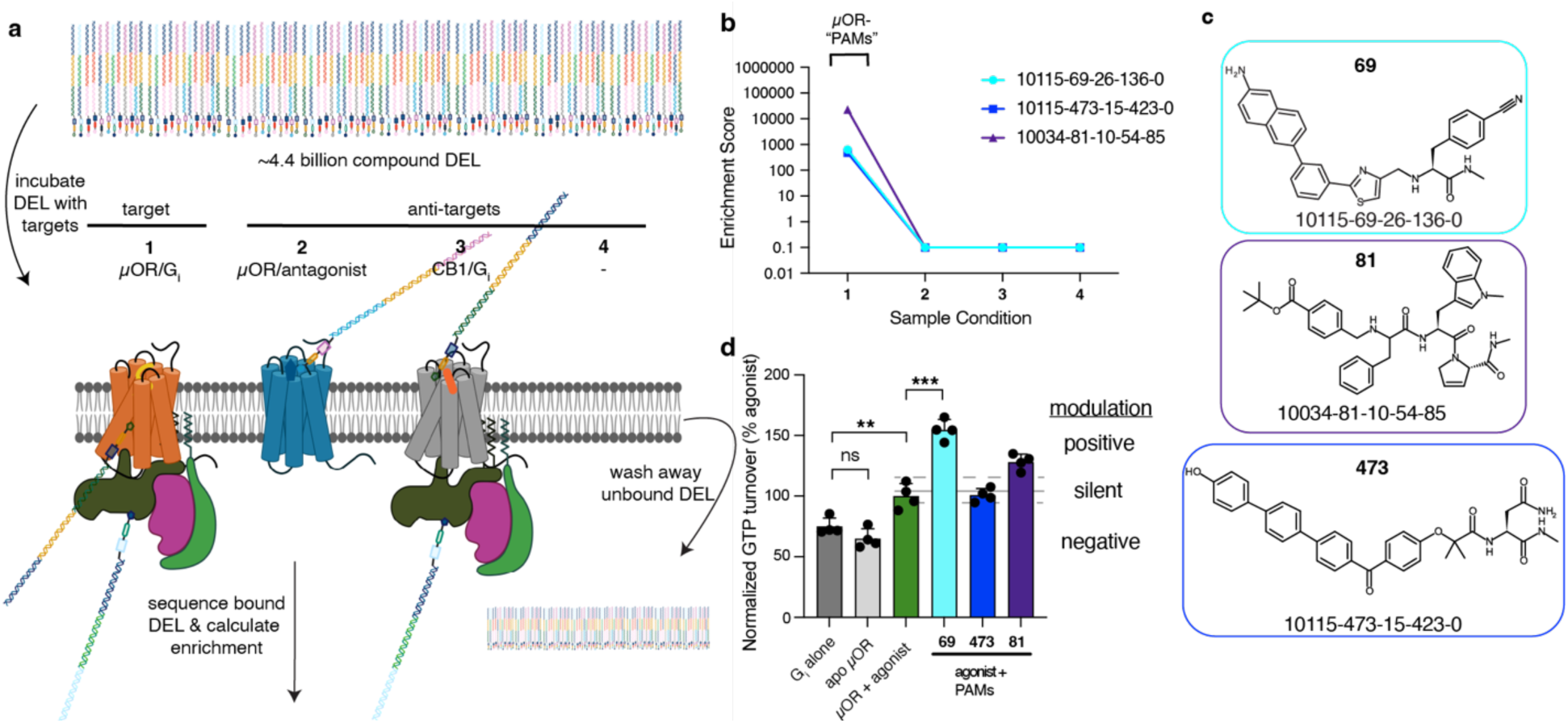
DEL screen for new μOR allosteric modulators. (**a**) Schematic detailing the DEL selection scheme including the active-state target (1 – µOR/G_i_,), and conditions used as anti-targets for removing inactive-state specific binders (2 – inactive µOR), G_i1_-specific binders (3 – CB1/G_i_), and non-specific binders (4 – no target control). Figure created with BioRender. Enrichment scores (**b**) and chemical structures (**c**) for the selected and characterized molecules across the 4 sample conditions. (**d**) GTP turnover assay used to initially assess the activity of the selected compounds for the µOR. G_i_ alone has intrinsic GTPase activity (dark gray) that is not significantly impacted by the presence of apo µOR (light gray) but is enhanced when the receptor is bound to agonist (green). PAMs for the µOR are expected to further enhance GTP turnover (i.e. **69**, 81), neutral silent modulators will have no effect and NAMs inhibit agonist-induced turnover.

## Results

### DNA-encoded library screening for GPCR PAMs

To identify PAMs that could enhance the analgesic effects of endogenous opioid peptides, we bound the µOR to met-enkephalin, one of the most abundant opioid peptides in the brain^18,19^, and then formed an active, nucleotide-free complex with a predominant G protein signaling partner of the µOR, G_i1_ (target **1, Fig. 1a**). Any DEL components enriched in this condition are likely to bind to the active ensemble of the receptor in a site distinct and non-competitive with enkephalin (and possibly other endogenous opioid peptides). Conversely, to identify (and remove) any molecules that bind to the inactive state of the receptor and may inhibit receptor activation (as in our previous study detailing the discovery and characterization of µOR NAMs^13^), we selected against DEL components that are enriched to µOR saturated with naloxone (anti-target **2, Fig. 1a**). These two conditions then serve as counter-selections for each other; selective enrichment in one and not the other indicates that a molecule is indeed sensitive to the activation state of the receptor. To remove compounds that bind to G protein rather than receptor, we counter-screened with active cannabinoid receptor 1 (CB1) bound to the potent full agonist MDMB-Fubinaca^20^ (FUB) and the same G_i1_ used for active µOR (anti-target **3, Fig. 1a**). Finally, no target control (anti-target **4, Fig. 1a**) eliminates molecules that bind non-specifically to the beads used to immobilize receptors.

After several rounds of binding the DEL to all 4 conditions independently and washing away unbound components, the enriched components were subjected to next-generation sequencing and relative enrichment scores were calculated based on the number of sequence reads in each condition (**Fig. 1b**). There were multiple “families” of enriched compounds with sequence counts observed *only* in the active state (condition **1**) of the µOR, with enrichment values (and raw sequence counts) ranging from moderate (634 enrichment, 8 sequence counts) to large (>20,000 enrichment, 24 sequence counts) (**Table S1**). From families of enriched compounds, we then selected the smallest, most polar representative compounds, including examples from three µOR PAM families (**Fig. 1b, 1c**). Compounds were then screened biochemically using an *in vitro* GTP-turnover assay^21,22^ where we assessed their ability to enhance agonist-dependent GTP depletion. Two of the three compounds selected as prospective µOR PAMs because of their specific enrichment to active µOR did indeed enhance met-enkephalin-induced activation of G protein (**Fig. 1d**), **81** and **69** (10115-69-26-136-0; calculated chemical properties in **Table S2**). **473** did not appear to modulate signaling of the receptor, though this may be due to either a lack of receptor binding or a lack of change in activation state upon binding. We chose to continue characterizing the mechanism of action of **69** because it was more efficacious (**Fig. 1d**) and potent (**Fig. S1b**) at improving GTP turnover compared to **81** while also having a smaller scaffold (**Table S2**). None of the molecules exerted these effects in the absence of the receptor (**Fig. S1c**).

Similar to our previous success in synthesizing a single “hit” that turned out to be a potent µOR NAM^13^, two out of three selected PAMs not only display binding to their target but modulate activity in the desired direction. Such high hit rates suggest that our steered selection methodology is a promising approach for efficient discovery of GPCR allosteric modulators.

### Pharmacological and biochemical assessment of **69** µOR PAM activity

Binding of **69** enhanced met-enkephalin-induced turnover with a roughly low double-digit micromolar potency as assessed by the GTP turnover assay (**Fig. 2a**). Though it has a low potency, it was highly effective at enhancing activity, fully depleting nucleotide under these conditions and converting met-enkephalin into a super-agonist. Consistent with its binding to (and presumably stabilizing) the active ensemble of µOR conformations, increasing amounts of **69** decreased the binding of ^3^H-naloxone to µOR-expressing membranes, with a comparable potency to that observed in the GTP turnover assay (**Fig. 2b**). This decrease in ^3^H-naloxone binding is likely due to a ∼6-fold decrease in naloxone affinity for the receptor in the presence of **69** (**Fig. S1d**). While binding of **69** decreased the binding of the orthosteric antagonist, it did not decrease binding of ^3^H-DAMGO, a full peptide reference agonist for the receptor (**Fig. 2c**). This is also consistent with the excess concentrations of peptide agonist (met-enkephalin) used in the selection conditions (**Fig. 1a**); such PAMs would not be predicted to be competitive with peptide agonists and indeed **69** is not. While increasing amounts of **69** do not decrease ^3^H-DAMGO binding (**Fig. 2c**), it does not enhance the agonist’s affinity^6^ or potency^23^ for the receptor like other µOR PAMs such as BMS-986122 or analogues, suggesting a different pharmacological mechanism of action.

**Figure 2.**
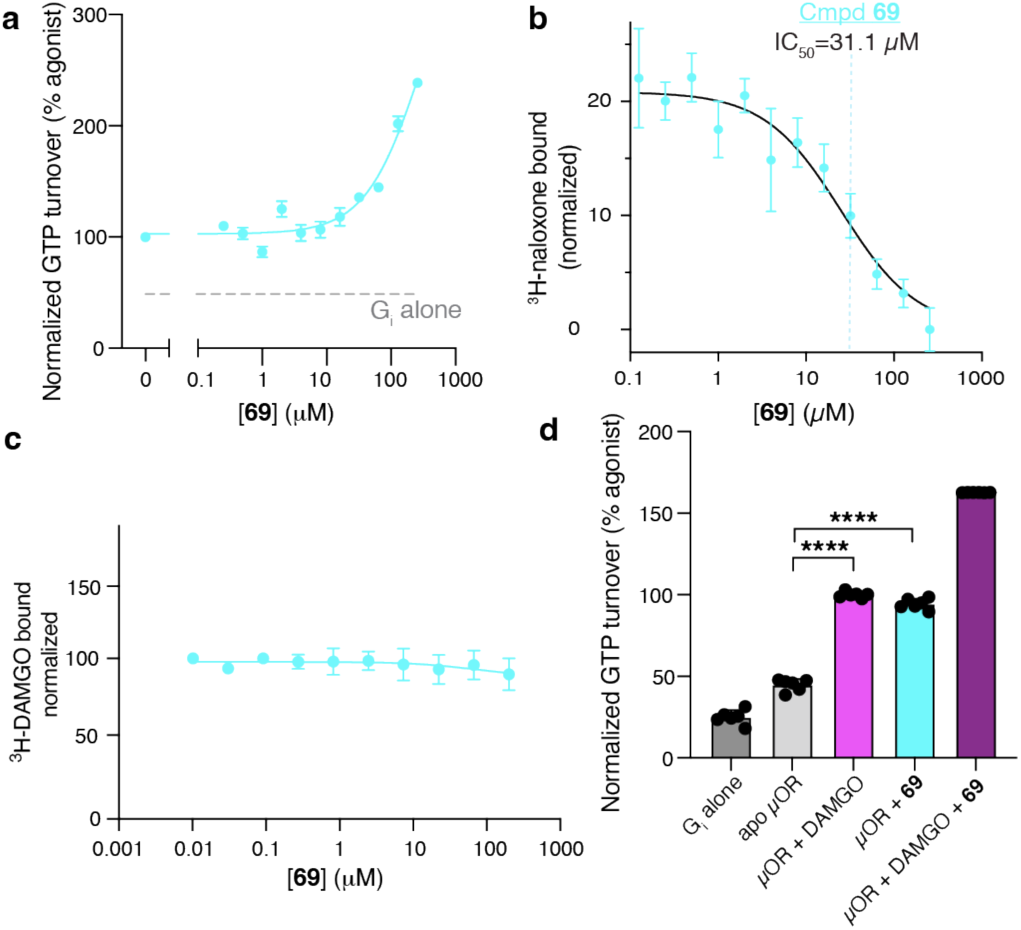
Biochemical activity of µOR PAM. (**a**) **69** was added in increasing amounts to the GTP turnover assay, demonstrating a significant increase in receptor-dependent activity but with a relatively low potency (∼30 µM). (**b**) “Competition” binding experiment showing that increasing concentrations of **69** result in decreasing amounts of bound antagonist (^3^H-naloxone), again with a double-digit µM affinity for the receptor. The observed IC_50_ value (31.1 µM, 95% CI of 14.4 to 38.8 µM) is displayed as a dashed cyan line. (**c**) ^3^H-DAMGO binding experiment demonstrating that while **69** does not have a substantial impact on agonist affinity for µOR. (**d**) GTP turnover assay with excess concentrations of **69** and/or full agonist peptide DAMGO showing that **69** alone can activate the receptor nearly to the same extent as DAMGO, and, consistently, further stimulates DAMGO-dependent turnover.

In addition to its effect on endogenous opioid peptides, **69** enhanced the efficacy of all orthosteric agonists tested, including the very weak partial/neutral antagonist naloxone and the partial agonist mitragynine pseudoindoxyl (MP)^24,25^, full agonist peptides (met-enkephalin, DAMGO), and a potent small molecule full agonist (BU72) (**Fig. S1e**). Further, in the absence of an orthosteric agonist, **69** alone induced GTP turnover to similar levels observed for the full agonist DAMGO, classifying it as an agonistic-PAM (ago-PAM) (**Fig. 2d**).

### 69 is active in membrane-based in vitro assays

We were next interested in characterizing how **69** impacts signaling at the cellular level. To begin to assess its impact on agonist-induced intracellular signaling cascades, we tested for cooperative activity with opioid agonists in the TRUPATH assay^26^, which measures G-protein heterotrimer dissociation upon activation of a GPCR. Here, we began by probing the ability of **69** to enhance DAMGO-induced dissociation of G_i1_ G-proteins. To our surprise, we did not see any significant impact on agonist-induced turnover (**Fig. 3a**). Given our clear biochemical (**Fig. 1d, 2a, 2d**) and membrane binding activity (**Fig. 2b, 2c**), we hypothesized that **69** may be binding to the interior region of µOR and is unable to cross the cell membrane. To test this hypothesis, we turned to the GTPγS binding assay, which measures binding of ^35^S-GTPγS to active G-proteins in *membranes*, rather than intact cells. Single-point versions of this assay demonstrate that, indeed, increasing concentrations of **69** enhance the efficacy of various concentrations of the orthosteric agonists buprenorphine (**Fig. 3b**) and methadone (**Fig. 3c**). Dose-response curves of the GTPγS binding assay show that increasing concentrations of **69** have minimal impact for the full agonist DAMGO (**Fig. 3d**) and substantial enhancements in buprenorphine potency and efficacy (**Fig. 3e**). This enhancement in orthosteric agonist efficacy is especially clear for the extremely weak partial agonist BP1-94, where no GTPγS binding is observed without our new PAM compound (**Fig. 3f**). This cooperativity with various orthosteric agonists is also seen in our biochemical GTPase assay (**Fig. S1e**). In summary, while **69** does not appear to cross the cell membrane, it is active in membrane-based assays and works in a generally cooperative manner with orthosteric agonists as in our initial biochemical assays.

**Figure 3.**
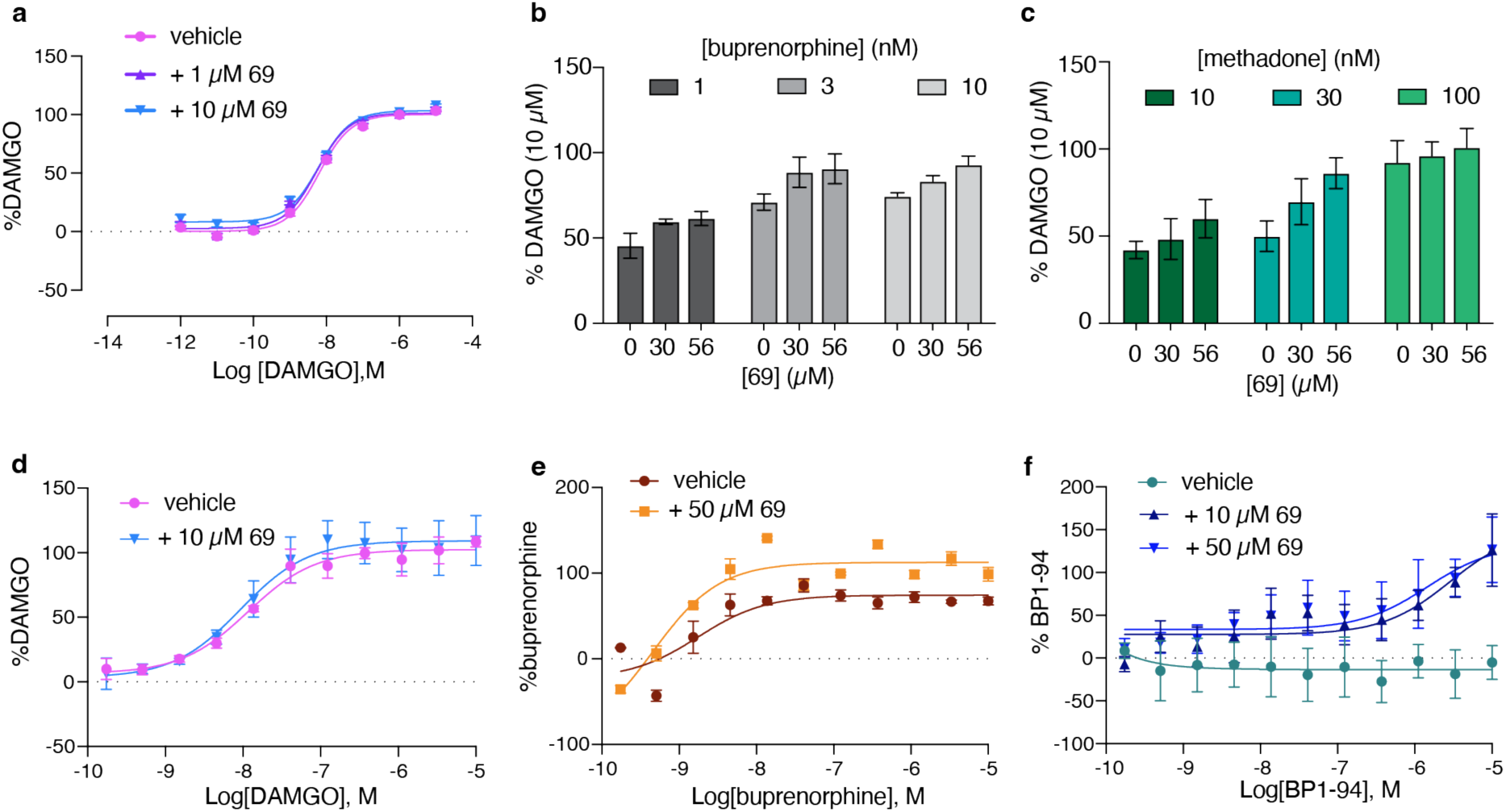
*69* enhances agonist efficacy in membranes but not cells. (**a**) G_i1_ TRUPATH experiment demonstrating that **69** has no substantial effect on DAMGO potency or efficacy in cell-based assays. Single-point GTPγS experiments demonstrate that increasing concentrations of **69** enhance the efficacy of various concentrations of the partial agonists buprenorphine (**b**) and methadone (**c**). This effect is further confirmed in dose-response versions of the GTPγS experiment where **69** has minimal impact on the full agonist DAMGO (**d**), but substantial increases in the efficacy of partial agonists like buprenorphine (**e**) and very weak near-antagonists like BP1-94 (**f**).

### Effect of PAMs on the dynamic activation behavior of µOR

GPCRs, including µOR, have extremely complex underlying energy landscapes that govern their multifaceted signaling outcomes^21,27–32^. Previous work using solution NMR has demonstrated the impact of µOR PAMs like BMS-986122 on the µOR activation landscape as well as provided clues about its binding site and mechanism of action^29^. Here, we aimed to use a direct readout of µOR conformational dynamics to better understand the role of allosteric modulators in influencing the outward movement of the key 6^th^ transmembrane helix (TM6). To do this, we applied a similar strategy to our previous single-molecule fluorescence resonance energy transfer (smFRET) study to investigate signaling bias in µOR^32^ where we labeled the receptor with a donor and acceptor fluorophore at this dynamic TM6 as well as a (largely static) reference position in TM4 (**Fig. 4a**). Labeled µOR was then attached to a surface via its N-terminal FLAG tag for long-term imaging of individual molecules. As in our previous work, in the presence of various orthosteric ligands alone, there is an efficacy-dependent change in FRET of a single major peak, with antagonist naloxone centered ∼0.90 and full agonist DAMGO centered ∼0.85 and partial agonist mitragynine pseudoindoxyl (MP) in between these values (**Fig. 4b** and **4d**, FRET histograms on left and fitted peak centers as bars on right). We interpret this as reflecting the fast exchange of TM6 between a closed **inactive** state and an **agonist-associated partially-open intermediate state**—a dynamic equilibrium subject to ligand-dependent regulation—which aligns with previous observations in both µOR^32^ and β2AR^21^. Agonist-associated intermediates have also been observed in the glucagon receptor, a Family B GPCR, though the rates of exchange between states in that system are much slower.^33^

**Figure 4.**
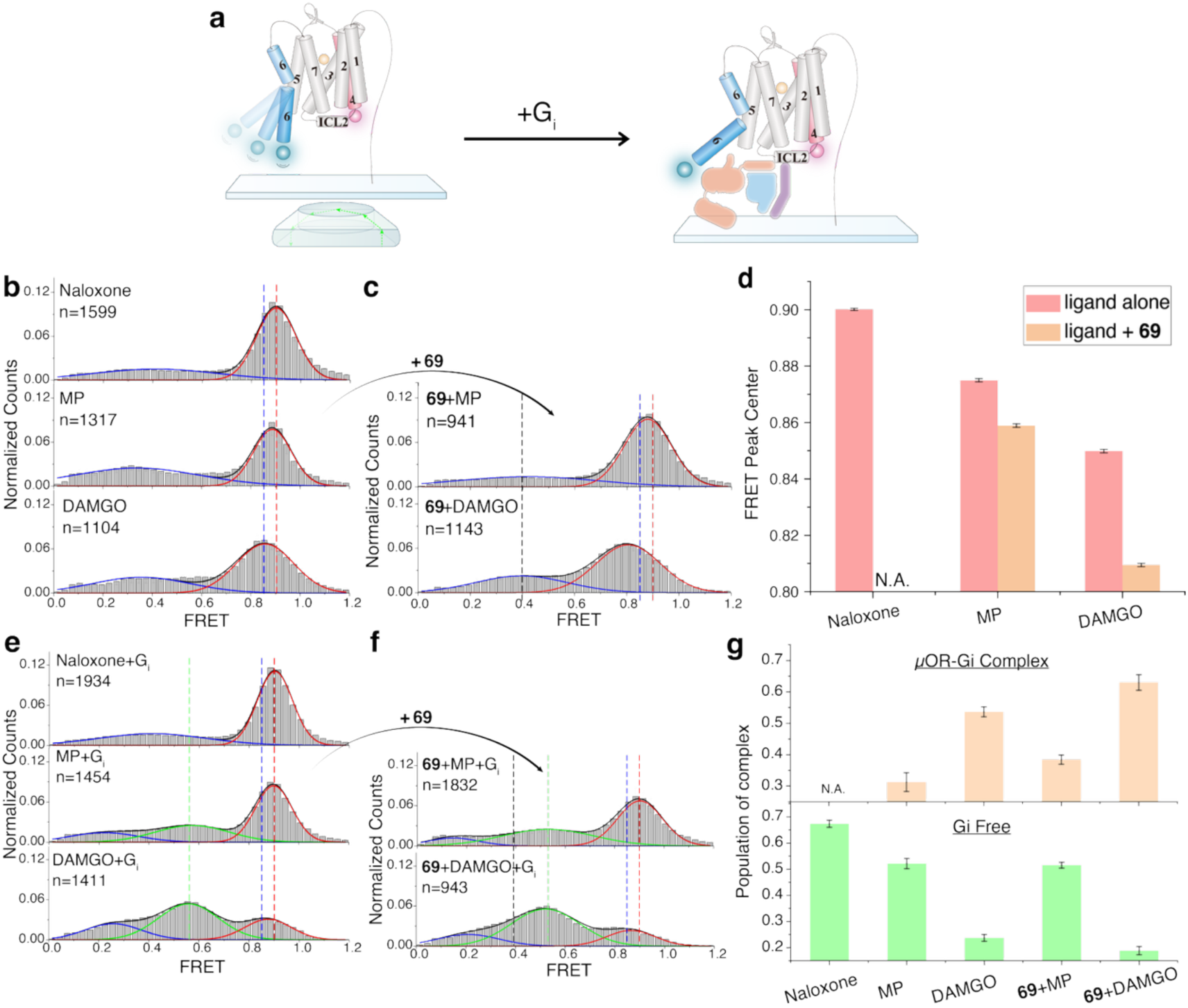
The effect of 69 on the conformational dynamics and structural changes of µOR. (**a**) Schematic of single-molecule FRET experiment and labeling sites. Labeled µOR was tethered to a cover slip via its Flag tag and biotinylated M1 Fab. (**b-c, e-f**) SmFRET distributions of µOR-IA-Cy3/Cy5 in the presence of different ligands without G_i_ (**b-c**) or with G_i_, followed by treatment of apyrase to remove GDP (**e-f**). Gaussian peaks were fitted to FRET states (red and green) and background noise (blue). Black lines represent the cumulative distributions. *n* represents the number of single-molecule traces used to calculate the corresponding histograms. The dash lines represent the FRET peak centers of different populations (Green: the G_i_-bound population centered ∼0.55; Blue/Red: the majority high-FRET states induced by DAMGO/Naloxone centered ∼0.85/∼0.90. Data are mean ± s.d. from three repeats. (**d**) FRET peak centers of µOR-IA-Cy3/Cy5 in the presence of different ligands w/o (pink bar) or w/ (brown bar) **69** in the absence of G_i_ extracted from the Gaussian fitting in (**b-c**). Standard errors are shown. (**g**) The population of µOR–G_i_ complex (orange bar) and G_i_-free μOR (green bar) in the presence of different ligands extracted from (**e-f**). Standard errors are shown.

Introduction of **69** to unliganded (apo) µOR results in a slight left-shifting in the high-FRET peak (peak value 0.84, **Fig. S2a**), though in the absence of orthosteric compounds, a predominant low-FRET state (∼0.39) emerges. As the observation of apo-state in our previous work^32^ and here (**Fig. S2a**), the distribution is dominated by broad, low-FRET signals, which likely arise from unstable/misfolded receptors. This makes interpretation of these low-FRET distributions challenging. The well-characterized µOR PAM compound BMS-986122, with distinct pharmacological properties compared with **69**, also results in high populations of this low-FRET state (**Fig. S2a**) in the absence of orthosteric compound. To exclude the possibility that **69** directly interacts with the fluorophores attached on µOR, we performed fluorescence anisotropy measurements. The fluorescence anisotropy (*r*) values of Cy5 remained essentially unchanged across five experimental conditions, including µOR in apo state and in the presence of DAMGO, naloxone, BMS-986122 or compound **69** (**Table S4**). The *r* values are close to or even smaller than 0.2, indicating that the rotation of the fluorophore is not significantly restricted and that the FRET efficiency is primarily determined by the distance between the fluorophores.^34^ Together, these results argue against the possibility that **69** strongly interacts with fluorophores to affect their rotational motions.

We next investigated whether **69** resulted in cooperative changes in TM6 conformation and intracellular G_i1_ binding with orthosteric agonists. While DAMGO binding causes a significant left-shift in the FRET center of the major FRET peak relative to the antagonist naloxone (**Fig. 4b**), addition of **69** to DAMGO-bound µOR results in a significant further decrease in the center of this peak to a FRET value of ∼0.81 (**Fig. 4d**). This is consistent with the allosteric nature of **69**, where its effects on TM6 are additive with orthosteric agonists (as in biochemical and membrane-based experiments in **Fig. 2** and **Fig. 3**, respectively). Addition of G_i1_ heterotrimers in the nucleotide-free state to agonist-bound µOR results in the selective population of a middle-FRET **active, G_i_-coupled state** centered ∼0.55 (**Fig. 4e, 4f**), presumably corresponding to the conformation observed in cryoEM structures of the G-protein bound receptor^29,35–37^. This peak is most prevalent for the full agonist DAMGO, where most of the receptors occupy this G_i1_-bound state with the remaining populations spread between the agonist-associated high-FRET peak and a low-FRET noise-associated peak. Unsurprisingly, little G-protein coupling state is observed when the receptor is bound to the antagonist naloxone (**Fig. 4e**). **69** alone does not appear to induce formation of the middle-FRET (*E*∼0.55) G_i_-coupled state on its own, nor does BMS-986122 (**Fig. S2b**). However, they do work cooperatively with agonists to enhance G_i_-engagement; DAMGO alone only results in ∼50% of the population in this G_i_-bound conformation, where the addition of **69** or BMS-986122 to DAMGO-bound receptors results in enhanced coupling with G_i1_ (**Fig. 4f, 4g, Fig. S2b, S2c**).

In summary, our new µOR PAM **69** can promote a leftward shift in the FRET efficiency of the agonist-associated intermediate state in a cooperative manner with orthosteric agonists, suggesting further outward movements of TM6, just like the already published PAM BMS-986122 (**Fig. S2b**). Lastly, although **69** alone appears insufficient to efficiently enhance G_i_ coupling, it still works cooperatively with DAMGO to ultimately enhance coupling with G_i_, consistent with the enhanced efficacy of this combination observed in both biochemical (**Fig. 2d**) and membrane-signaling (**Fig. 3**) assays.

## Discussion

We^13^ (and others^14,15^) have shown that we can use DEL hits as useful biochemical and biophysical tools to investigate mechanisms of activation and allostery in GPCRs. We’ve also shown that these hits can act as therapeutic leads to treat opioid overdose and substance use disorder^13^. Here, we develop and implement a further advance in the DEL selection methodology to selectively screen for functional positive (and negative) allosteric modulators of GPCRs by counter-screening active, agonist and G-protein bound states of the receptor against inactive, antagonist-bound states. Control selections can then be used to select against G-protein-specific and non-specific binders. Discovery of GPCR allosteric modulators is historically quite challenging with traditional methods; this strategy will hopefully serve as a generalizable method for streamlined discovery of such compounds.

After demonstrating the functionality of 2/3 of the prospective PAMs synthesized off-DNA, we used a combination of biochemical, pharmacological, and cell/membrane-based signaling experiments to characterize the mechanism of action of our lead compound, **69**. This µOR PAM enhanced the efficacy of a wide array of orthosteric agonists in biochemical and membrane-based signaling assays, but is apparently not functional in intact cell-based assays. This leads us to speculate that **69** has a binding site on µOR either on the intracellular side of the receptor or within the membrane and cannot efficiently cross the lipid bilayer.

Finally, we employed smFRET to directly visualize how µOR PAMs modulate the conformational dynamics of the key 6^th^ transmembrane helix (TM6), thereby elucidating the conformational landscape of activation. Our new PAM works cooperatively with DAMGO to further stabilize its “conventional” agonist-associated intermediate state, directly resulting in enhanced interaction with G_i_ partner protein (**Fig. 4e, 4f, 4g**). This cooperativity is likely underlying the observed enhancement in efficacy in both biochemical and membrane-based efficacy assays (**Fig. 2, Fig. 3**). Future work will seek to establish the binding site of **69** on µOR using structural biology to confirm its intracellular or intra-membrane binding site speculated here, while further medicinal chemistry will seek to enhance the membrane permeability of the compound and enhance its potency and efficacy as a µOR PAM. Such molecules will be candidates to enhance endogenous opioid antinociception, resulting in pain relief with mitigated opioidinduced side effects.^4^

## Acknowledgements

We thank WuXi Apptec for providing the DEL as well as J. Su for providing SAR information for chemical families. K.K. was supported by the American Diabetes Association (ADA) Postdoctoral Fellowship. E.S.O. was supported by the American Heart Association (AHA) Postdoctoral Fellowship. B.K.K. was supported by the Chan Zuckerberg Biohub and by NIH R01DA036246. C.C. was supported by the National Natural Science Foundation of China Grant No. 22425701 and the National Key R&D Program of China Grant No. 2024YFA0916700 and 2024YFA1306200. J.M.S. and P.T. were supported by NIH R01DA052340. S.M. is supported by DA059978 from National Institute of Health.

## Author Contributions

E.S.O., & B.K.K. conceived the study and wrote the manuscript with input from all authors. E.S.O, K.K., W.H. & B.K.K. designed the DEL screening strategy. E.S.O. performed DEL selections with counter-target provided by K.K. E.W. and E.S.O. designed, optimized, and performed radioligand binding experiments. E.S.O. and Y.S. optimized and performed GTP turnover assays. J.W. performed smFRET experiments under the supervision of C.C.M.L. performed single-point GTPγS experiments under the supervision of J.R.T. P.T. performed titration versions of the GTPγS experiments under the supervision of J.M.S. B.P. and K.A. performed cell-signaling experiments under the supervision of S.M.

## Methods

### Purification of µ-opioid receptor and variants

#### Purification of mouse µ-opioid receptor from sf9 cells

The mouse µ-opioid receptor was grown and purified as previously described^35^. Mouse µ-opioid receptor (µOR) with an N-terminal FLAG and C-terminal hexa-histidine tag was expressed as previously described^35^ using the baculovirus method in Spodoptera frugiperda (Sf9) cells. Naloxone was added to 10 µM final concentration upon infection and cells were collected 48 hours post-infection and stored at-80°C for later purification. µOR was extracted from membranes with 0.8% *n*-dodecyl-ß-D-malopyranoside (DDM; Anatrace), 0.08% cholesterol hemisuccinate (CHS), and 0.3% 3-((3-cholamidopropyl) dimethylammonio)-1-propanesulfonate (CHAPS; Anatrace) in 20 mM HEPES pH 7.5, 500 mM sodium chloride (NaCl), 30% glycerol, 5 mM imidazole, 10 µM naloxone, and the protease inhibitors benzamidine and leupeptin, along with benzonase (Sigma-Aldrich) to degrade cellular DNA. Cells were dounced 30 times on ice and the membranes were allowed to solubilize in the detergent for 2 hours with stirring at 4 °C, followed by centrifugation for 40 minutes at 14k rpm to pellet cell debris. The supernatant was applied to nickel-chelating sepharose resin and bound to resin with end-over-end shaking for 2 hours at 4°C. The resin was then batch washed 4 times followed by washing with 10 column volumes on-column with nickel wash buffer composed of 20 mM HEPES pH 7.4, 500 mM NaCl, 0.1% DDM, 0.01% CHS, 0.03% CHAPS, 5 mM imidazole, 10 µM naloxone, and protease inhibitors leupeptin and benzamidine. The nickel-pure µOR was then eluted in the same buffer with 250 mM imidazole. The nickel elution was initially exchanged to lauryl maltose neopentyl glycol (L-MNG; Anatrace) detergent by incubating with 0.5% L-MNG, 0.17% glycol-diosgenin (GDN; Anatrace) and 0.067% CHS overnight at 4°C. 2 mM calcium chloride (CaCl_2_) was then added and subsequently applied to M1 anti-FLAG immunoaffinity resin. The M1-bound receptor was first washed with 20 mM HEPES pH 7.4, 500 mM NaCl, 0.1% MNG, 0.033% GDN, 0.0133% CHS, 2 mM CaCl_2_, and 10 µM naloxone. The protein was then washed with 10 column volumes of 20 mM HEPES pH 7.4, 100 mM NaCl, 0.005% MNG, 0.0017% GDN, 0.00067% CHS and 2 mM CaCl_2_ followed by elution with 20 mM HEPES pH 7.4, 100 mM NaCl, 0.003% MNG, 0.001% GDN, 0.0004% CHS, 5 mM ethylenediaminetetraacetic acid (EDTA) and FLAG peptide. Multimers and dimers of the receptor were removed with size exclusion chromatography on an S200 10/300 Increase gel filtration column (GE Healthcare) equilibrated with 20 mM HEPES pH 7.4, 100 mM NaCl, 0.01% MNG, 0.001% CHS. Pure, monomeric apo µOR was spin concentrated to ∼150 µM, flash frozen in liquid nitrogen and stored at-80°C until further use. This receptor was used for DEL selections and cryoEM studies.

#### Purification of human µ-opioid receptor from Expi293 cells

The gene for the full-length human µ-opioid receptor was cloned into a vector for inducible expression in Expi293F cells (Thermo Fisher) with N-terminal HA signal peptide and FLAG tags and a C-terminal hexahistidine tag. This construct was transfected into Expi293F cells constitutively expressing the tetracycline repressor (Thermo Fisher) with the Expifectamine transfection kit (Thermo Fisher) following manufacturers directions with induction of receptor expression 2 days after transfection with doxycycline (4 µg/mL and 5 mM sodium butyrate) in the presence of 10 µM naloxone. Pellets were collected and frozen at-80 °C ∼30 hours after induction for later protein purification. Cells were dounced and membranes solubilized with 20 mM HEPES pH 7.5, 100 mM NaCl, 20% glycerol, 1% L-MNG, 0.1% CHS, 10 µM naloxone, protease inhibitors leupeptin and benzamidine, and benzonase, followed by purification on anti-FLAG immunoaffinity resin as above. Multimers and dimers of the receptor were removed with size exclusion chromatography on an S200 10/300 Increase gel filtration column (GE Healthcare) equilibrated with 20 mM HEPES pH 7.4, 100 mM NaCl, 0.01% MNG, 0.001% CHS. Pure, monomeric apo µOR was spin concentrated to ∼150 µM, flash frozen in liquid nitrogen and stored at-80°C until further use. This receptor was used in parallel with the above mouse µOR to assess compound activity against human µOR in GTPase assays.

### Purification of cannabinoid receptor 1

Purification of human full-length CB1 with an N-terminal FLAG and C-terminal hexahistidine tags was performed as previously described using the baculovirus method (Expression Systems).^20^ Briefly, CB1 was extracted from insect cell membranes with 1% L-MNG and 0.1% CHS and purified by nickel-chelating sepharose chromatography. Crude, nickel-pure CB1 was applied to M1 anti-FLAG immunoaffinity resin and washed with decreasing amounts of inverse agonist SR. The receptor was eluted from M1 resin with a buffer consisting of 20 mM HEPES pH 7.5, 150 mM NaCl, 0.05% L-MNG, 0.005% CHS, FLAG peptides and 5 mM EDTA.

The FLAG-pure CB1 was further purified by size exclusion chromatography on an S200 10/300 Increase gel filtration column (GE Healthcare) equilibrated with 20 mM HEPES pH 7.5, 150 mM NaCl, 0.05% L-MNG and 0.005% CHS. Purified, monomeric CB1 was concentrated to ∼500 µM and flash frozen and stored at-80°C until further use.

### Expression & purification of heterotrimeric G-proteins

Heterotrimeric G_i_ was expressed in *Trichoplusia ni* (*T. ni*) with the BestBac method (Expression Systems) and purified as previously described.^20,35^ Briefly, T. ni cells were infected with one virus encoding the wild-type human Gα_i_ subunit and another encoding the wild-type human β_1_ γ _2_ subunits with a histidine tag inserted at the N-terminus of the β subunit. Cells were harvested 48 hours post infection and lysed with hypotonic buffer. Heterotrimeric Gα_i_ β_1_ ©_2_ proteins were extracted in a buffer containing 1% sodium cholate and 0.05% DDM. The heterotrimer was purified using nickel-chelating sepharose chromatography while removing cholate. Human rhinovirus 3C protease (3C protease) was added to cleave off the histidine tag overnight at 4°C on-column. The flow through was collected and dephosphorylated with lambda protein phosphatase (NEB), calf intestinal phosphatase (NEB) and Antarctic phosphatase (NEB) in the presence of 1 mM manganese chloride (MnCl_2_). The heterotrimer was further purified by ion exchange chromatography on a MonoQ 10/100 GL column (GE Healthcare) in 20 mM HEPES pH 7.5, 1 mM MgCl_2_, 0.05% DDM, 100 µM TCEP, and 20 µM GDP and eluted with a linear NaCl gradient from 50 to 500 mM. The purified heterotrimer was collected and dialyzed into 20 mM HEPES pH 7.5, 100 mM NaCl, 0.02% DDM, 100 µM TCEP and 20 µM GDP overnight at 4°C, followed by concentration to <250 µM, addition of 20% glycerol and flash-freezing in liquid nitrogen and storage at-80°C until further use.

### Preparation of samples for DEL selections

#### µOR/G_i_/met-enkephalin complex formation

µOR/G_i_ complex was formed in a similar manner to that described previously^35,36^. Briefly, excess met-enkephalin peptide [MedChemExpress] was added to purified µOR (above) and incubated at room temperature for 1 hour. Concurrently, 1% L-MNG/0.1% CHS was added to G_i_ purified in DDM to exchange detergent on ice for 1 hour. The two reactions were mixed for a final molar ratio of 1:1.5 µOR:G_i_ and incubated at room temperature for 1 hour. Apyrase was added and the reaction was further incubated on ice for 2 hours. 2 mM CaCl_2_ was added to the reaction before adding to M1 anti-FLAG resin for 20 minutes. The column was washed with 20 mM HEPES pH 7.4, 100 mM NaCl, 0.003% MNG, 0.001% GDN, 0.0004% CHS, 5 µM met-enkephalin, and 2 mM CaCl_2_, followed by elution in the same buffer with 5 mM EDTA replacing the CaCl_2_ and FLAG peptides. After removing excess unbound G_i_ heterotrimer, excess un-coupled µOR was removed by size exclusion chromatography with a S200 10/300 Increase column into 20 mM HEPES pH 7.4 and 100 mM NaCl, 0.003% MNG, 0.001% GDN, 0.0004% CHS, 100 µM TCEP and 1 µM met-enkephalin. µOR/G_i_/met-enkephalin complex was spin concentrated to ∼25 µM and flash-frozen in liquid nitrogen after adding 15% glycerol.

#### MOR/naloxone formation

Purified µOR (above) was incubated with excess naloxone for 1 hour at room temperature before selection experiments in 20 mM HEPES pH 7.4, 100 mM NaCl, 0.003% MNG, 0.001% GDN, 0.0004% CHS.

#### CB1/G_i_/MDMB-fubinaca complex formation

CB1/G_i_ complex was formed in a similar manner to that described previously^20^. Briefly, excess MDMB-fubinaca agonist was added to purified CB1 (above) and incubated at room temperature for 1 hour. Concurrently, 1% L-MNG/0.1% CHS was added to G_i_ purified in DDM to exchange detergent on ice for 1 hour. The two reactions were mixed for a final molar ratio of 1:1.25 CB1:G_i_ and incubated at room temperature for 3 hours. Apyrase was added and the reaction was further incubated on ice for 1.5 hours. 2 mM CaCl_2_ was added to the reaction before adding to M1 anti-FLAG resin for 20 minutes. The column was washed with 20 mM HEPES pH 7.4, 100 mM NaCl, 0.003% MNG, 0.001% GDN, 0.0004% CHS, 5 µM MDMB-fubinaca, and 2 mM CaCl_2_, followed by elution in the same buffer with 5 mM EDTA replacing the CaCl_2_ and FLAG peptides. After removing excess unbound G_i_ heterotrimer using M1 anti-FLAG immunoaffinity chromatography, excess un-coupled CB1 was removed by size exclusion chromatography with a S200 10/300 Increase column into 20 mM HEPES pH 7.4 and 100 mM NaCl, 0.003% MNG, 0.001% GDN, 0.0004% CHS, 100 µM TCEP and 10 µM MDMB-fubinaca. CB1/G_i_/MDMB-fubinaca complex was spin concentrated to ∼25 µM and flash-frozen in liquid nitrogen.

### DEL selection

#### Target preparation

The three samples produced above (µOR/G_i_/met-enkephalin, µOR/naloxone, and CB1/G_i_/MDMB-fubinaca) were diluted to 2 µM final concentration in **W1**: 20 mM HEPES pH 7.4, 150 mM NaCl, 100 µM TCEP, 0.1% hsDNA (Thermo Fisher Scientific), 0.02% MNG, 0.002% CHS and 20 µM ligand (met-enkephalin, naloxone, and MDMB-fubinaca, respectively). A fourth blank sample (no protein control) was made with the same buffer in the absence of ligand. 300 µL of HisPur^TM^ Ni-NTA magnetic beads (Thermo Fisher Scientific) as slurry were washed with water followed by splitting into 4 batches and washing with the 4 respective buffers made above. Washing with buffer was repeated 2 additional times. 300 µL of 2 µM targets (above) and 300 µL buffer without agonist was incubated with the respective magnetic Ni-NTA resin for 30 minutes with mixing to bind targets to resin. Each of the 4 samples was then split into 3 100 µL aliquots corresponding to each of the 3 rounds of selection.

#### First round of selection

Any unbound target was removed from the 4 100 µL targets by washing with 200 µL respective wash buffer above. The 4 DEL samples (G1, G2, G3, G4) provided by WuXi (10 µL) were resuspended with 90 µL of the respective wash buffers above for the 4 conditions and then applied to the 4 targets bound to magnetic resin and incubated with shaking at room temperature for 1.5 hours to bind to the target(s). Reactions were spun briefly to settle the resin, followed by washing all with 200 µL respective buffer above. The 4 selections were then washed again with 200 µL of the same buffers with lower detergent concentrations (0.001% MNG, 0.0001% CHS) and lower ligand concentrations (500 nM) (**W2**). The 4 selections were then washed with 200 µL of the same respective buffers in the absence of detergent and even further lower ligand concentration (100 nM) (**W3**) to minimize detergent and ligand carry-over between rounds. The resin with target (and bound library molecules) was then resuspended in 100 µL of the respective last wash buffer and heated at 95 °C for 10 minutes to elute bound library components but leave receptor bound to resin. The heat denatured mixture was spun down and the supernatant removed from the resin. 10 µL of each of the 4 selections was saved for later analysis, and 90 µL was reserved for the next round of selection.

#### Second round of selection

Another 100 µL aliquot of target-bound magnetic resin was washed with 200 µL respective **W1** buffer to remove unbound target. The 4 samples of resin were then resuspended in 90 µL respective **W1**, to which the 90 µL of reserved Round 1 DEL selection was added and incubated with shaking at room temperature for 1.5 hours. Reactions were spun briefly to settle the resin, followed by washing with 200 µL respective **W1** buffer. The 4 selections were then washed again with 200 µL of respective **W2** buffer, followed by 200 µL of respective **W3** buffer. The resin with target (and bound library molecules) was then resuspended in 100 µL respective **W3** and heated at 95 °C for 10 minutes to elute bound library components but leave receptor bound to resin. The heat denatured mixture was spun down and the supernatant removed from the resin. 50 µL of each of the 4 selections was saved for later analysis, and 50 µL was reserved for the next round of selection.

#### Third round of selection

Another 100 µL aliquot of target-bound magnetic resin was washed with 200 µL respective **W1** buffer to remove unbound target. The 4 samples of resin were then resuspended in 50 µL respective **W1**, to which the 50 µL of reserved Round 2 DEL selection was added and incubated with shaking at room temperature for 1.5 hours. Reactions were spun briefly to settle the resin, followed by washing with 200 µL respective **W1** buffer. The 4 selections were then washed again with 200 µL of respective **W2** buffer, followed by 200 µL of respective **W3** buffer. The resin with target (and bound library molecules) was then resuspended in 100 µL respective **W3** and heated at 95 °C for 10 minutes to elute bound library components but leave receptor bound to resin. The heat denatured mixture was spun down and the supernatant removed from the resin. All of the final (third) round of selection supernatant for all 4 conditions was saved for subsequent sequencing and enrichment analysis.

### Preparation of µ-opioid receptor membranes

Cell membranes containing the mouse µOR described above were generated by infecting Sf9 cells in an identical manner to that described above for protein purification. Sf9 cells expressing µOR were resuspended in cold lysis buffer composed of 10 mM HEPES pH 7.4, 10 mM MgCl_2_, and 20 mM potassium chloride (KCl) with the protease inhibitors leupeptin, benzamidine, and cOmplete^TM^ EDTA-free protease inhibitor cocktail tablets (Sigma-Aldrich). The lysed cells were spun at 45k rpm for 45 minutes to pellet membranes. The supernatant was removed and membrane pellets were resuspended in cold lysis buffer followed by douncing ∼30 times on ice. The dounced membranes were spun again at 45k rpm for 45 minutes. The pellets were resuspended in cold lysis buffer again with the addition of benzonase and dounced a further∼30 times, followed by a further spin at 45k rpm for 45 minutes. Membranes were resuspended in the same lysis buffer above in the presence of 1 M NaCl and dounced a further ∼30 times and pelleted. Finally, membranes were washed with the original lysis buffer, dounced, and pelleted. Final washed membranes were resuspended to 2g original pellet mass per 1 mL in lysis buffer; the resulting µOR-containing membranes were flash frozen for later radioligand binding experiments.

### Radioligand binding experiments

#### Membrane binding experiments

µOR-containing insect cell membranes prepared above were diluted 1:1000 in 20 mM HEPES pH 7.4, 100 mM NaCl, and 0.05% bovine serum albumin (BSA). For saturation binding experiments, membranes were incubated with a serial dilution of ^3^H-naloxone (50.3 Ci/mmol; Perkin Elmer) and allosteric modulators at different constant concentrations for 1.5 hours at room temperature with shaking. For “competition” binding experiments, membranes were incubated with 2 nM ^3^H-naloxone and serially diluted allosteric modulators for 1.5 hours at room temperature with shaking. Following incubation, membranes were rapidly bound to double thick 90 x 120 mm glass fibre Printed Filtermat B filters (Perkin Elmer) and washed with cold binding buffer (20 mM HEPES pH 7.4, 100 mM NaCl) using a MicroBeta Filtermat-96 cell harvester (Perkin Elmer). ^3^H-naloxone bound membranes on Filtermats were measured with a MicroBeta counter (Perkin Elmer) after addition of MultiLex B/HS melt-on scintillator sheets (Perkin Elmer) and data values were plotted after normalizing total counts per minute to the highest and lowest values.

### GTP turnover assay

The GTP turnover assay was performed using a modified version of the GTPase-GLO^TM^ assay (Promega) as described previously^21,22^. Purified µOR was diluted to 1 µM in 20 mM HEPES pH 7.4, 100 mM NaCl, 0.01% L-MNG, 0.001% CHS, and 20 µM guanosine-5’-triphosphate (GTP) in the presence of various orthosteric (20 µM met-enkephalin, 20 µM naloxone, 20 µM MP, 20 µM H-Tyr-D-Ala-Gly-*N*(Me)Phe-Gly-OH [DAMGO], 20 µM BU72) and allosteric (serially diluted or at excess concentrations of 100 µM for 10115-69-26-136) ligands and incubated for 1.5 hours at room temperature. Concurrently, G_i_ purified in DDM was exchanged by incubating with 1% L-MNG and 0.1% CHS for 1 hour on ice. The exchanged G_i_ was then diluted to 1 µM in 20 mM HEPES pH 7.4, 100 mM NaCl, 0.01% L-MNG, 0.001% CHS, 20 µM guanosine-5’-diphosphate (GDP), 200 µM TCEP, and 20 mM MgCl_2_. Equal volumes of receptor solutions and G_i_ solution were mixed and incubated at room temperature for 60 minutes (agonist-bound receptor experiments) or 90 minutes (apo receptor experiments) with gentle shaking. Controls include mixing equal volumes of both buffers (total initial GTP) and equal volumes of 1 µM G_i_ solution and receptor buffer (intrinsic G protein turnover). Equal volume of GTPase-Glo reagent supplemented with 10 µM adenosine 5’-diphosphate (ADP) in 20 mM HEPES pH 7.4, 100 mM NaCl, 0.01% MNG and 0.001% CHS was added and incubated with gentle shaking for 30 minutes, followed by addition of further equal volume of detection reagent. After brief (10 minute) incubation, luminescence was measured with a MicroBeta counter.

### PAM-mode TRUPATH (BRET) assays

G-protein dissociation was measured using the TRUPATH kit^26^ gifted by Bryan Roth (Addgene kit #1000000163) using experiments in PAM mode, similar to those previously described^7,9,38^. HEK293T (ATCC CRL-11268) cells were cultured in Dulbecco’s Modified Eagle Medium (DMEM) high glucose with GlutaMAX™ Supplement (Gibco™, 10566024) and 10% Fetal Bovine Serum (FBS) (GemCell™, 100-500). The cells were plated in TC-Treated Culture dishes (100mm x 20mm) (Corning™, 430167) at a seeding density of 2.5 million cells per dish and incubated overnight at 37°C. The next day, cells were transfected with a 1:5 DNA ratio of human Mu opioid receptor (µOR) to triple Gi1 containing inserts Gα-RLuc8, Gβ3, and Gγ9-GFP2 (AddGene Plasmid #196048). Lipofectamine 2000 (Invitrogen™, 11668019) was used to complex the DNA following the vendor’s protocol in Opti-MEM (Gibco™, 31985070), and incubation continued for 16 hours. The following day, cells were harvested using Trypsin-EDTA (0.05%) (Gibco™, 25300120) and plated in poly-D-lysine-coated (Sigma Aldrich P0899) 384-well assay plates (Greiner Bio-One, 781098) with white, clear bottoms at a density of 16,000–20,000 cells per well in DMEM with 1% dialyzed FBS (Gibco™, A3382001). The next day, the cell medium was decanted, a white backing (Brightmax™, Z732117) was applied to the plate bottom, and the cells were washed with 30 μL of assay buffer (1X Hank’s balanced salt solution (HBSS) (Gibco™, 14025092) + 20 mM HEPES, pH 7.4 (Gibco™, 15630080)). For PAM screening, 30 μL of different concentrations of compound 69 (0.1, 1, 10 μM) in assay buffer with 0.3% bovine serum albumin were added and incubated for 1 hour. Afterwards, the cells were treated with 15 μL of dose response for µOR agonist control, DAMGO (prepared as 4× drug concentration in assay buffer with 0.3% bovine serum albumin), followed by the addition of 15 μL substrate buffer (40 μM coelenterazine 400a (Cayman Chemical, 16157). This was incubated for 5 minutes in the dark at room temperature before reading. The plates were read every 5 minutes for four intervals using a BioTek Synergy Neo Alpha plate reader (Agilent Technologies, USA) with 395 nm (for RLuc8-coelenterazine 400a) and 510 nm (for GFP2) emission filters. Measurements from the 10-minute read were used for analyses. The ratio of GFP2/RLuc8 for G-protein was calculated per well in quadruplicates, plotted as a function of drug concentration, normalized to %DAMGO control agonist stimulation, and analyzed using “log(agonist) vs. response (three parameters)” in GraphPad Prism 10.2.1.

### GTPγS single-point experiments

#### Membrane Preparation

As previously described^23^ CHO cells stably expressing wild-type human μOR (hMOR-CHO) were grown to confluence at 37 °C in 5 % CO_2_ in Dulbecco’s modified Eagle media (DMEM) containing 10 % fetal bovine serum, 1 % penicillin/streptomycin and 400 ug/mL G418. Cells were detached from the plates by incubation in harvesting buffer (20 mM HEPES, pH 7.4, 150 mM NaCl, and 0.68 mM EDTA) at room temperature, and pelleted by centrifugation at 1000 g for 3 min. The cell pellet was suspended in ice-cold 50 mM Tris-HCl buffer, pH 7.4, and homogenized with a Tissue Tearor (Biospec Products, Inc.) for 20-30 s. The homogenate was re-centrifuged at 20,000 g for 20 min at 4 °C, and the pellet resuspended in 50 mM Tris-HCl, pH 7.4, followed by re-centrifugation. The final pellet was resuspended in 50 mM Tris-HCl, pH 7.4, to 0.5-1.0 mg/mL protein and stored frozen in aliquots at-80 °C. Protein concentration was determined using Bicinchoninic Acid Assay with bovine serum albumin as the standard.

### 35S-GTPγS assay binding assay

Membrane homogenates (20 μg protein) were incubated for 60 min in a shaking water bath at 25 °C in a buffer containing 50 mM Tris-HCl, pH 7.4, 5 mM MgCl_2_, 100 mM NaCl, 30μM GDP, 0.1 nM [^35^S]GTPγS, with 69 or vehicle in the presence of varying concentrations of buprenorphine or methadone as described^23^. Samples were filtered through the glass-fiber filters mounted in a Brandel harvester and rinsed five times with ice-cold 50 mM Tris-HCl, pH 7.4, containing 5 mM MgCl_2_, and 100 mM NaCl. Filters were dried, and following the addition of EcoLume scintillation cocktail, counted in a Wallac 1450 MicroBeta Liquid Scintillation and Luminescence Counter (Perkin Elmer). DAMGO as a control was included in all assay plates. The level of [^35^S]GTPγS bound is expressed as percentage of the response to the standard MOR agonist DAMGO (10 μM DAMGO).

### GTPγS concentration/response experiments

#### Cell culture and membrane preparation

A Chinese Hamster Ovary (CHO-K1) parental cell line expressing the human µOR was obtained from PerkinElmer (#ES-542-C) and used for all experiments. Cells were maintained in 1:1 DMEM/F12 media with 1x penicillin/streptomycin supplement and 10% heat-inactivated fetal bovine serum (all from Invitrogen/ThermoFisher) in a 37°C/5% CO_2_ incubator. Maintenance cultures were further supplemented with 500 μg/mL of G418 to preserve receptor selection/expression (Invitrogen/ThermoFisher). Cells were generally passaged at 1:10 every 2 days. Cells were collected with 5 mM EDTA in dPBS and pelleted by centrifuging at 4000 rpm for 10 minutes at 4°C. Pellets were stored at-80°C until use.

The pellet was resuspended in ice-cold homogenization buffer (10 mM Tris-HCl pH 7.4, 100 mM NaCl, 1 mM EDTA) and homogenized for three 10-second intervals at the maximum setting using a Polytron homogenizer, with 30-second cooling periods on ice between each burst. The homogenates were centrifuged (600g, 10 minutes, 4°C), the pellet was discarded, and the supernatant was re-centrifuged (20,000g, 1 hour, 4°C). The final pellet was resuspended in 20 mM HEPES pH 7.4, 10 mM MgCl_2_, and 100 mM NaCl using a syringe. The protein concentration was determined by using a BCA assay quantification method (Bio-Rad) with bovine serum albumin as the standard. The resulting aliquots were further stored at –80°C until required for the assay.

### 35S-GTPγS assay

The ^35^S-GTPγS coupling assay was performed as in our previous work^39,40^. Briefly, 20 μg of cell membrane protein was combined with 0.124 nM of ^35^S-GTPγS (PerkinElmer, #NEG030H250UC) and concentration curves of ligands (see the figure legends for details) in a 200 μL reaction volume in the presence of 40 μM GDP. The reactions were incubated at RT for 1 hr; they were then collected onto GF/C filter plates (PerkinElmer) using a Brandel Cell Harvester. Then, the plates were read on a PerkinElmer Microbeta2 scintillation counter. The resulting data were normalized to stimulation caused by vehicle (0%) and reference agonist (100%) and fit to a 3-variable (Hill Slope = 1) agonist model using GraphPad Prism 10.0.

### Single-molecule FRET experiments

#### µOR preparation and labeling

Methods for sample preparation and single-molecule measurements were adapted from our previous work.^32^ A minimal-cysteine µOR construct (µORΔ7), in which all solvent-exposed cysteines were mutated (C13S, C22S, C43S, C57S, C170T, C346A, C351L), was used. Then, two labeling sites, R182C and R273C, located on TM4 and TM6, respectively, were subsequently introduced.

The receptor was expressed in Sf9 insect cells (authenticated by the supplier and not tested for mycoplasma) using the Bac-to-Bac baculovirus expression system and purified by size-exclusion chromatography using a Superdex 200 Increase 10/300 column (GE Healthcare), following previously established procedures. Purified µOR was labeled with homemade iodoacetamide-conjugated sulfo-Cy3 and sulfo-Cy5 fluorophores (Lumiprobe) as described previously.^32^ Excess dye was removed using a home-packed desalting column containing 2 ml G50 resin (Sigma) equilibrated with desalting buffer (20 mM HEPES, pH 7.5, 100 mM NaCl, 0.01% LMNG, 0.001% CHS, 15% glycerol). The final concentration of labeled receptor was approximately 1 μM.

#### Single-molecule FRET experiments and analysis

All single-molecule experiments were performed at 25 °C using a custom-built objective-type total internal reflection fluorescence (TIRF) microscope based on a Nikon Ti-E platform. A 532 nm solid-state laser (OBIS Smart Lasers, Coherent Inc.) was used for excitation, and fluorescence emission was detected using an EMCCD camera (Andor iXon Ultra 897). Emission signals were separated using a dichroic mirror (T635lpxr, Chroma) and bandpass filters (ET585/65m for Cy3 and ET700/75m for Cy5) mounted in a Dual-View spectral splitter (Photometrics). Instrument control and data acquisition were performed using Cell Vision software (Beijing Coolight Technology). smFRET movies were acquired at 100 ms/frame with a laser power of 10 mW.

Fluorophore-labeled µOR was immobilized on polyethylene glycol (PEG) passivated glass coverslips via biotinylated M1-Fab and streptavidin. Glass chambers were cleaned and passivated with the mixture of PEG and biotin–PEG, followed by incubation with 0.05 mg/ml streptavidin in buffer containing 20 mM HEPES (pH 7.5) and 100 mM NaCl. After 1 min, unbound streptavidin was removed by washing with incubation buffer (50 mM HEPES, pH 7.5, 100 mM NaCl, 0.01% LMNG, 0.001% CHS, 2 mM CaCl₂, 5 mM MgCl₂, and 100 μM ligand/PAM). Biotinylated M1-Fab (10 nM) was then introduced in incubation buffer and incubated for 1 min, followed by removal of unbound Fab. ∼ 20 nM N-terminal FLAG-tagged, fluorophore-labeled µOR was pre-incubated with ligands (w/ or w/o PAMs) on ice for 20 min, subsequently diluted to ∼1 nM, and introduced into the chamber. Unbound receptor was removed by washing with imaging buffer consisting of incubation buffer supplemented with an oxygen scavenging and triplet-state quenching system (50 nM protocatechuate-3,4-dioxygenase (PCD), 2.5 mM protocatechuic acid (PCA), 1.5 mM Trolox, 1 mM 4-nitrobenzyl alcohol (NBA), and 1 mM cyclooctatetraene (COT)). For measurements involving nucleotide-free G_i1_, µOR (20 nM) pre-incubated with 100 μM ligand was mixed with 20 μM G_i1_ and incubated for 20 min. Apyrase (New England Biolabs) was then added at a 1:100 (v/v) ratio to deplete residual nucleotides. After an additional 1 h incubation on ice, the complex was diluted, introduced into the imaging chamber, and measured following the same protocol described above. Where indicated, 100 μM BMS-986122 or PAM69 was included together with orthosteric ligands, except conditions containing the antagonist naloxone.

Single-molecule fluorescence trajectories were extracted from recorded movies using a custom-written ImageJ plugin. Fluorescence spots were identified and fitted with a two-dimensional Gaussian function, and donor–acceptor spot pairs were matched using a variant of the Hough transform. Background-subtracted integrated intensities of the fitted Gaussian peaks were used as the raw fluorescence intensities. Apparent FRET efficiency (*E*) was calculated as: 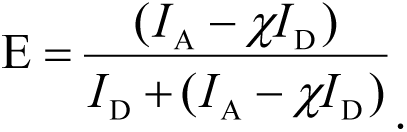 where *I*_D_ and *I*_A_ denote the raw donor and acceptor fluorescence intensities, respectively, and χ represents donor bleed-through into the acceptor channel (χ = 0.05). FRET traces were selected using a custom MATLAB script based on two criteria^41^: (1) a signal-to-noise ratio greater than 4, defined as the mean total intensity prior to photobleaching divided by its standard deviation; and (2) the presence of a single-step photobleaching event in the donor channel. Frame-wise FRET efficiencies prior to photobleaching were pooled to generate FRET efficiency histograms, which were fitted using Gaussian functions.

### Fluorescence anisotropy assay

All fluorescence anisotropy measurements were performed at 25 °C using a commercial multimode plate reader Envision-I (PerkinElmer). The optical configuration and acquisition settings for Cy5 fluorescence anisotropy were established according to the manufacturer’s instructions, and the G-factor was calibrated prior to data collection.

Receptor incubation conditions were identical to those used in single-molecule experiments performed in the presence of ligand alone. Following incubation, receptor samples were transferred to black 96-well plates (OptiPlate^TM^-96F, PerkinElmer), for 100 µL per well and fluorescence polarization values (*P*) were recorded. Fluorescence anisotropy (*r*) was calculated from the measured polarization values according to the following equation: 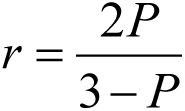

### Figure Preparation

Graphs were created using *GraphPad Prism.* Schematic figures were created using *BioRender*. Figures were constructed in *Adobe Illustrator*.

## Supplementary Figures

**Figure S1.**
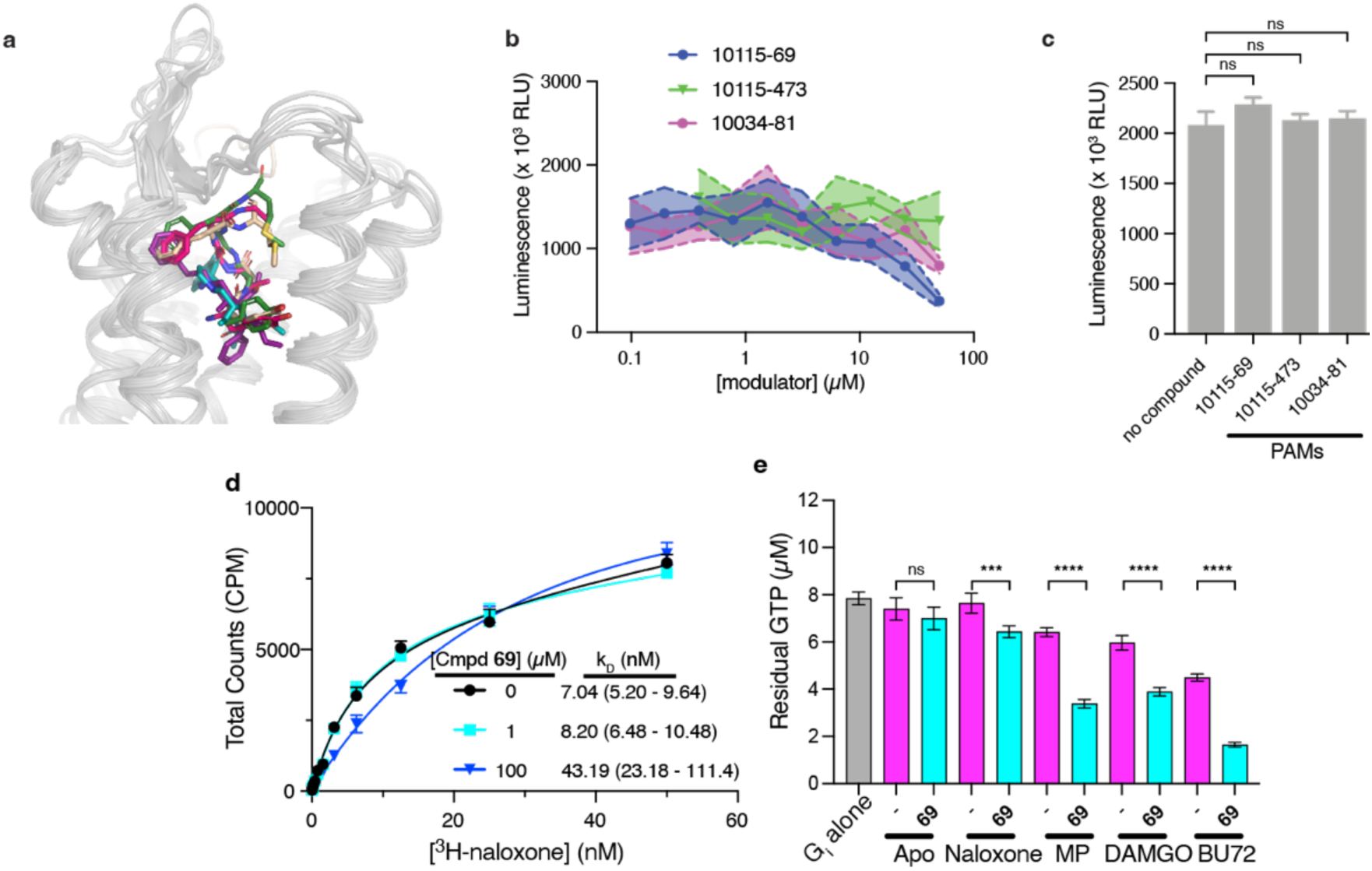
Initial biochemical characterization of µOR allosteric modulators from DEL screen. (**a**) Alignment of several active, G-protein bound structures of the µOR bound to orthosteric agonists, ranging including small molecules and peptides. The protein backbone is colored grey and agonists displayed as sticks (met-enkephalin, green, present work; β-endorphin, wheat, PDB: 8F7Q; DAMGO, magenta, PDB: 6DDE; mitragynine pseudoindoxyl, teal, PDB: 7T2G; lofentanil, purple, PDB: 7T2H). (**b**) The GTP turnover assay was used to assess rough compound potency for their ability to modulate agonist-bound µOR-induced G_i1_ turnover. Titration of the 3 selected µOR-specific potential modulators from the DEL screen demonstrates that **69** and **81** enhance receptor activity with double-digit µM potency. Error bars correspond to standard deviations of four measurements. (**c**) Excess concentrations of all µOR-specific potential modulators have no impact on G_i1_ intrinsic turnover in the absence of receptor. Error bars correspond to standard deviations of four measurements. Differences between all conditions relative to no compound control are not significant (P > 0.05) using an unpaired t-test. (**d**) Using a direct ^3^H-naloxone binding experiment, we show that excess concentrations of **69** result in a decreased antagonist affinity for µOR-containing membranes. Fitted affinity values are shown along with 95% confidence intervals. Error bars represent the standard deviations of four measurements. (**e**) The GTP turnover assay was used to show that **69** enhances turnover for a wide variety of orthosteric site conditions, ranging from weak (naloxone) to moderate (mitragynine pseudoindoxyl, MP) partial agonists to peptide (DAMGO) or small molecule (BU72) full agonists.

**Figure S2.**
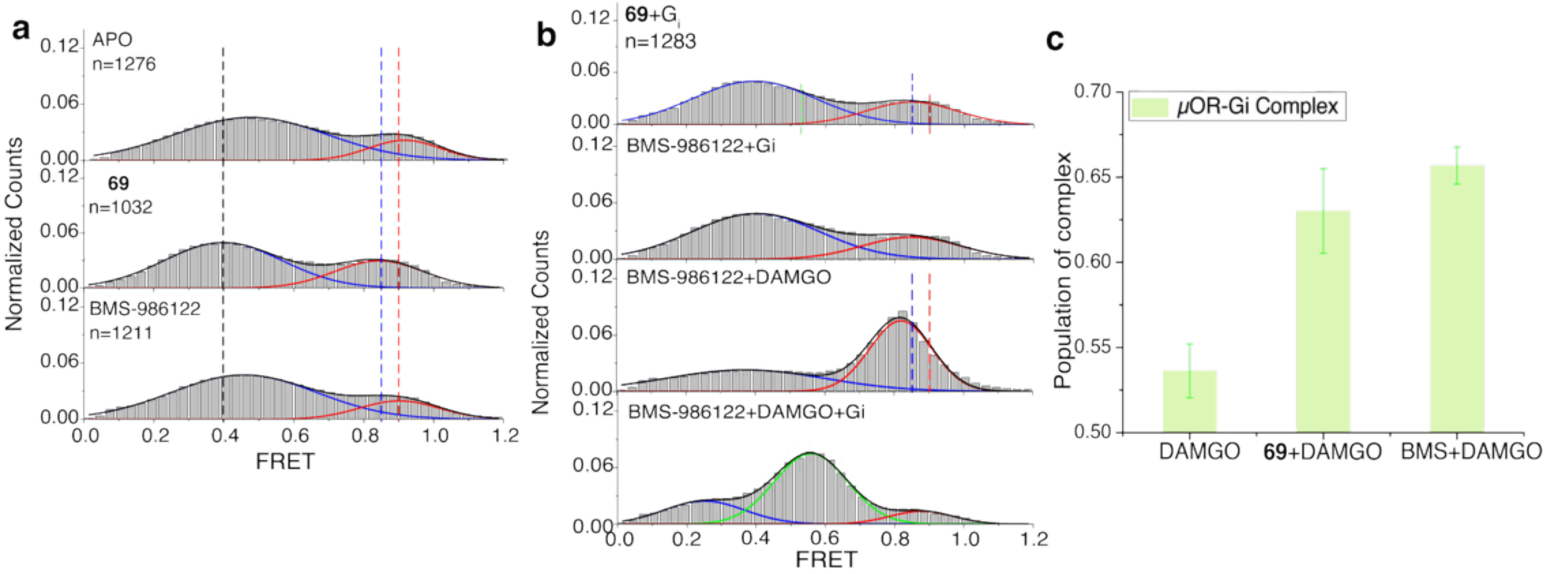
Novel TM6 conformational state detection for PAM-bound opioid receptor. (**a-b**) SmFRET distributions of µOR-IA-Cy3/Cy5 in apo state or in the presence of different ligands and PAMs without (**a**) or with (**b**) G_i_. Gaussian peaks were fitted to FRET states (red and green) and background noise (blue). Black lines represent the cumulative distributions. *n* represents the number of fluorescence traces used to calculate the corresponding histograms. The dash lines represent the FRET peak centers of different populations. Black: the low-FRET peak observed in the absence of orthosteric ligand at ∼0.39; Blue/Red: the majority high-FRET states induced by DAMGO/Naloxone centered ∼0.85/∼0.90, respectively. Data are mean ± s.d. from three repeats. (**c**) The population of µOR–G_i_ complex in the presence of DAMGO with/without PAMs. Standard errors are shown.

## Supplementary Tables

**Table S1.** Enrichment of compounds in various selection conditions.

| Compound | Counts C1 | Counts C2 | Counts C3 | Counts C4 | Enrichment C1 | Enrichment C2 | Enrichment C3 | Enrichment C4 |
| --- | --- | --- | --- | --- | --- | --- | --- | --- |
| 10115-69-26-136-0 | 24 | 0 | 0 | 0 | 634.5 | 0.1 | 0.1 | 0.1 |
| 10115-473-15-423-0 | 18 | 0 | 0 | 0 | 475.87 | 0.1 | 0.1 | 0.1 |
| 10034-81-10-54-85 | 8 | 0 | 0 | 0 | 22789.94 | 0.1 | 0.1 | 0.1 |

**Table S2.**
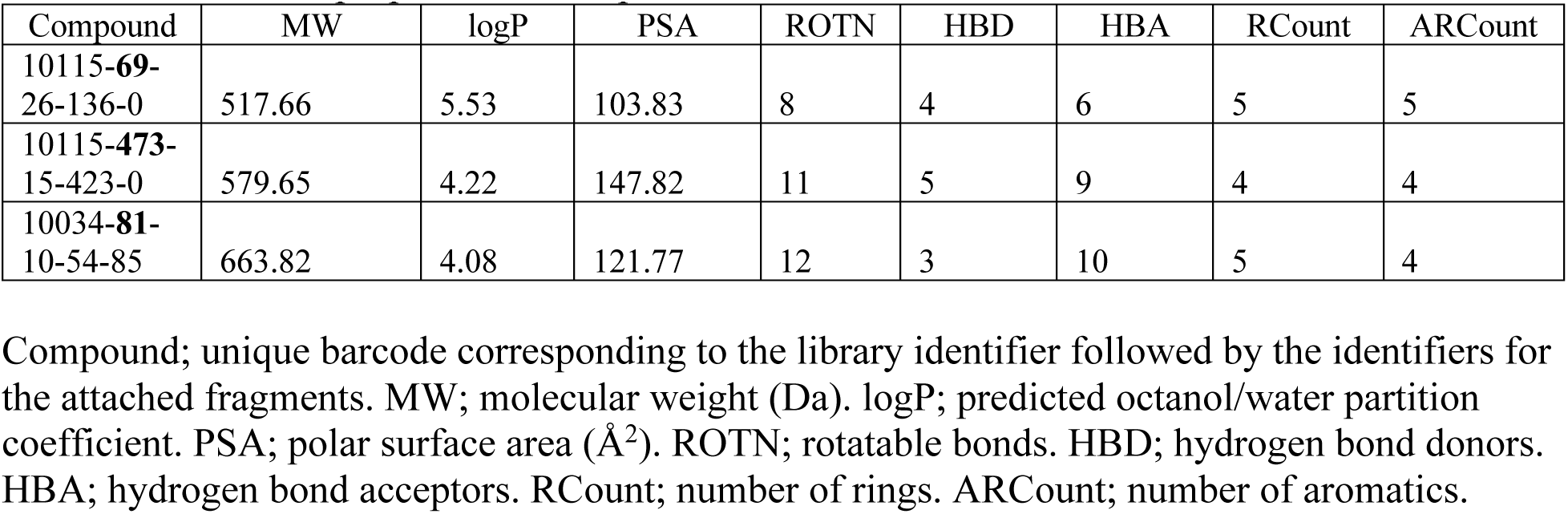
Chemical properties of compounds.

**Table S3.** Enrichment properties of selected chemical families.

| Compound | Counts<br>C1 | Counts<br>C2 | Counts<br>C3 | Counts<br>C4 |
| --- | --- | --- | --- | --- |
| Cmpd **69** |  |  |  |  |
| 10115-69-26-136-0* | 24 | 0 | 0 | 0 |
| 10115-69-12-136-0 | 10 | 0 | 0 | 0 |
| 10115-185-16-136-0 | 7 | 0 | 0 | 0 |
| 10115-290-7-136-0 | 7 | 0 | 0 | 0 |
| 10115-69-15-423-0 | 6 | 0 | 0 | 0 |

Sequence counts in each of the four selection conditions for family members selected for further study (asterisks identify the exact molecule selected for synthesis and characterization). For **69**, fragment 1 (identifier 69) and fragment 3 (identifier 136) appear to be important for binding to condition 1, while fragment 2 is more variable.

**Table S4.** Fluorescence anisotropy (*r*) of Cy5 labeled on μOR under different conditions.

| Incubation conditions | APO | DAMGO | Naloxone | BMS-986122 | Cmpd **69** |
| --- | --- | --- | --- | --- | --- |
| $r \pm$ s.d. | 0.192±0.004 | 0.198±0.006 | 0.197±0.005 | 0.194±0.003 | 0.192±0.006 |

The fluorescence anisotropy ( $r$ ) of Cy5 fluorophore labeled on μOR in apo state and in the presence of DAMGO, naloxone, BMS-986122 or compound **69**. Data are mean ± s.d. from three repeats.

